# Silencing of lncRNA NEAT1 inhibits esophageal squamous cell carcinoma proliferation, migration, and invasion via regulation of the miR-1299/MMP2 axis

**DOI:** 10.1101/2021.01.19.427233

**Authors:** Zhanfeng Yang, Meilin Zhu, Jingjing Zhang, Guangchong Zhang, Jun Hu, Qunli He

**Affiliations:** Department of Medicine, Zhengzhou University of Industrial Technology, Xinzheng, Henan, China; School of Basic Medical Sciences, Zhengzhou University, Zhengzhou, Henan, China; Henan Medical College, Zhengzhou, Henan, China

**Keywords:** lncRNA NEAT1, Esophageal squamous cell carcinoma, MiR-1299, CeRNA, MMP2

## Abstract

Esophageal squamous cell carcinoma (ESCC) is the most prevalent form of esophageal cancer worldwide. Considerable evidence has verified that abnormal expression of lncRNAs can effectively influence the progression of various malignant tumors. However, the regulatory mechanisms of lncRNAs underlying ESCC development and progression remain poorly defined. Here, we investigated the role of lncRNA nuclear-enriched abundant transcript 1 (NEAT1) in ESCC via regulating microRNA 1299 (miR-1299) and matrix metalloproteinase 2 (MMP2). A total of 32 ESCC tissue samples were obtained from the First Affiliated Hospital of Zhengzhou University. The mRNA levels of lncRNA NEAT1, miR-1299, and MMP2 mRNA were measured via quantitative real-time PCR. Interactions among miR-1299, lncRNA NEAT1, and MMP2 mRNA in EC9706 cells were confirmed by dual-luciferase reporter assays and RNA immunoprecipitation (RIP) assays. The proliferation and migration/invasion of ESCC cells were verified by CCK-8 and transwell assays, respectively. lncRNA NEAT1 was up-regulated in ESCC tissues and cells. lncRNA NEAT1 silencing inhibited migration, invasion, and proliferation of ESCC cells. Furthermore, lncRNA NEAT1 sponged and negatively regulated miR-1299, thus giving rise to increased expression of MMP2. Moreover, miR-1299 inhibitors and MMP2 rescued the invasion of ESCC cells following silencing of lncRNA NEAT1. lncRNA NEAT1 was overexpressed in ESCC tissues and cells. Silencing of lncRNA NEAT1 inhibited ESCC proliferation, migration, and invasion via reducing competitive binding of lncRNA NEAT1 with miR-1299 and enhancing miR-1299-targeted suppression of MMP2. Taken together, our findings suggest that lncRNA NEAT1 is a potential target for ESCC therapy and rehabilitation.

---

Esophageal squamous cell carcinoma (ESCC) is the most prevalent form of esophageal cancer worldwide and exhibits a high incidence rate and fatality rate worldwide, especially in East Asia, East Africa, South Africa, and Southern Europe[1, 2]. Clinically, despite enormous advances having been made in preoperative radiotherapy, chemotherapy, and surgery in the past several years, the chance of survival for ESCC patients has been improved but the five-year survival rate has remained at only 15%–20%, with a median survival time of approximately 1.5 years[3]. Moreover, because the early symptoms are not obvious, the overwhelming majority of ESCC patients are diagnosed at an advanced stage when tumors cannot be thoroughly surgically removed[4]. Poor prognosis is a direct result of neoplasm metastasis, radiotherapy resistance, and the high recurrence rate of ESCC.[5] Thus, identifying more exploitable molecular markers for early and metaphase diagnosis, as well as elucidating underlying regulatory mechanisms of ESCC, are urgently needed for ESCC patients to improve prognosis and survival. Long non-coding RNAs (lncRNAs), each of which are over 200 nt in length, are transcribed by RNA polymerase and are produced by splicing and modification of small nuclear or mature RNAs, but do not have any capacity for encoding proteins[6]. Previous evidence has indicated that lncRNAs may regulate numerous biological processes such as tumor growth and metastasis, including those involved in ESCC[7]. Moreover, lncRNAs play roles as signals, scaffolds, guides, decoys, and competing endogenous RNAs (ceRNAs) through interacting with their corresponding target genes[8]. In various cancers, many lncRNAs have been discovered to exhibit abnormal expression levels and to be closely related to tumor progression. For instance, lncRNA ZFAS1 is up-regulated in ESCC and promotes cell proliferation, migration, and invasion in ESCC cell lines by downregulating miR-124 and upregulating STAT3[9]. lncRNA KLF3-AS1 is down-regulated in ESCC and inhibits tumor growth and invasion via weakening miR-185-5p-mediated inhibition of KLF3[10]. lncRNA PANDA is upregulated in ESCC and drifts away from NF-YA to promote the expression of NF-YA-E2F1 co-regulated proliferation-promoting genes, as well as to restrict apoptosis[11]. lncRNA HAGLR is highly expressed in esophagus cancer (EC), and HAGLR depletion leads to cell proliferation, metastasis, invasion, and inhibition of the epithelial-mesenchymal transition (EMT) by restoring the expression of miR-143-5p and downregulating LAMP3[12]. Additionally, nuclear enriched abundant transcript 1 (NEAT1) is an lncRNA that has been verified to be a carcinogen in diverse malignancies, including colorectal cancer, breast cancer, hepatocellular carcinoma, lung cancer, melanoma cancer, prostate cancer, pancreatic cancer, and ovarian cancer (OC)[13-20].

Furthermore, many studies have demonstrated that abnormal expression patterns of miRNAs serve as tumor-inhibiting factors and carcinogens in multiple human malignancies, including ESCC[21]. MiR-1299 has been shown to be down-regulated in diverse cancers, including colon cancer, prostate cancer, hepatocellular carcinoma, and ovarian cancer; furthermore, miR-1299 has also been demonstrated to act as a tumor suppressor[22-25]. Additionally, matrix metalloproteinase 2 (MMP2) has been identified as an important gene for invasion and lymph-node metastasis in ESCC, and overexpression of MMP2 protein in ESCC tissues verifies that it is closely associated with esophageal metastasis[26]. According to the gene sequences of these aforementioned molecules matched via bioinformatic analysis and literature review, we hypothesized that lncRNA NEAT1 may influence proliferation, migration, and invasion of ESCC via regulating miR-1299 and MMP2. Hence, in the current study, we investigated the potential molecular mechanisms of lncRNA NEAT1 in regulating miR-1299 and MMP2 in ESCC, with the aim of identifying a novel strategy for ESCC therapy and rehabilitation.

## Materials and methods

### Main experimental materials

Normal human esophageal epithelial cells (Het-1A) and human ESCC cell lines (KYSE30, KYSE150, and EC9706 cells) were all provided by the Shanghai Cell Bank at the Chinese Academy of Sciences (Shanghai, China). SiRNA lncRNA NEAT1 (si-NEAT1), si-RNA negative controls (si-NCs), an miR-1299 inhibitors, miR-1299 mimics (miR-1299), and microRNA negative controls (miR-NCs) were obtained from RiboBio (Guangzhou, China). Lacking 3’-UTR MMP2 overexpression plasmids (pcDNA-MMP2) were purchased from GenePharma (Shanghai, China).

### Patient specimens

A total of 32 pairs of ESCC tissue and para-cancer tissue samples were obtained from the First Affiliated Hospital of Zhengzhou University. These patients had not received radiotherapy or chemotherapy before surgery. Immediately after surgical excision, these tissue specimens were stored directly in liquid nitrogen until further use. Ethical approval for this study was approved by the Ethics Committee of Zhengzhou University. All procedures were in accordance with the Declaration of Helsinki and written informed consent was obtained from each patient in advance.

### Cell cultures and transfections

Cells were cultured in RPMI-1640 (Hyclone, Logan, UT, USA) supplemented with 10% fetal bovine serum (FBS; Gibco, Grand Island, NY, USA) in a humidified atmosphere at 37°C with 5% (v/v) CO_2_. Si-NEAT1, si-NCs, miR-1299 mimics, miR-NCs, miR-1299 inhibitors, and pcDNA-MMP2 were transfected into KYSE30 and EC9706 cells via Lipofectamine 2000 reagent (Invitrogen, Carlsbad, CA, USA) according to the manufacturer’s instructions.

### Quantitative real-time PCR (qRT-PCR)

Total RNAs were extracted separately from tissue samples and cultured cells using TRIzol reagent (Invitrogen, Carlsbad, CA, USA) on the basis of the manufacturer’s protocol. Then, lncRNA NEAT1 and MMP2 mRNA were reversed transcribed into cDNA using PrimeScript RT Reagent Kit (Takara, Dalian, China), and miR-1299 was reversed transcribed into cDNA using TaqMan microRNA Reverse Transcription Kit (Invitrogen, Carlsbad, CA, USA). Subsequently, RNA levels were measured via quantitative real-time qPCR (qRT-PCR) on a Bio-Rad CFX96 Touch System (Bio-Rad, Hercules, CA, USA). The primers used in this study were synthesized by Tsingke (Tsingke, Beijing, China) and are listed in Table 1.

**Table 1.**
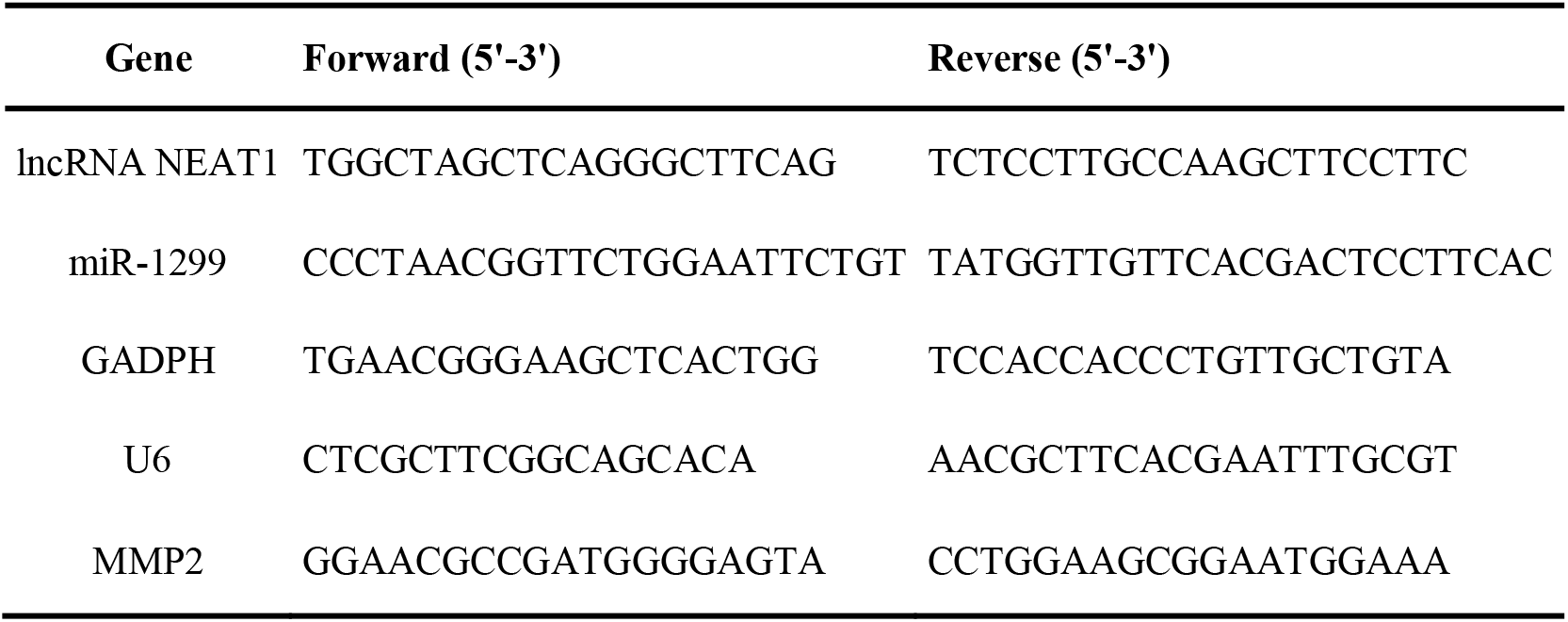
Primer sequences used for qRT-PCR

### Cell counting kit-8 (CCK8) assays

Cell proliferation was measured by Cell Counting Kit-8 assays (Dojindo Molecular Laboratories, Japan). Initially, the transfected KYSE30 and EC9706 cells were seeded in 96-well plates (2×10^3^ cells/well) and cultured for several days at 37°C with 5% (v/v) CO_2_. Then, 10 µL of CCK-8 solution was added to the cells and incubated for 1 h at 0, 24, 48, and 72 h. Subsequently, the optical density (OD) value of each sample at 450 nm was measured by a microplate reader (ELx800; BioTek, Winooski, VT, USA).

### Transwell migration and invasion assays

Transwell assays were conducted to assess the migration and invasion of ESCC cells. After transfection, KYSE30 and EC9706 cells (1×10^4^ cells/well) were suspended in serum-free medium and seeded onto the upper chamber of inserted 8-μm-pore filters coated with Matrigel (BD Biosciences, San Jose, CA, USA) for invasion assays or uncoated for migration assays. RPMI-1640 medium containing 10% FBS was added to the lower chamber. After incubation for 24 h, the cells having no invasive or migratory activities were erased from the upper surface of the chamber with cottons swabs and the invaded and migrated cells on the lower surface of the membrane were fixed with 4% paraformaldehyde for 30 min, after which they were stained with 0.3% crystal violet for 5 min. Then, nine regions were randomly selected and the numbers of cells in these regions were counted with an inverted microscope (Nikon, Tokyo, Japan).

### Dual-luciferase reporter assays

Bioinformatic analyses of miRDB and lncBase v2 were explored to search and obtain three binding sites of lncRNA NEAT1 and miR-1299. Then, pmirGLO plasmids containing three pairs of wild-type and mutant binding sequences were purchased from GeneChem (Shanghai, China), including pmirGLO-Wt1, pmirGLO-Mt1, pmirGLO-Wt2, pmirGLO-Mt2, pmirGLO-Wt3, and pmirGLO-Mt3. Meanwhile, wild-type and mutant MMP2 3′-UTRs were synthesized and recombined into pmirGLO Luciferase vectors (Promega, Madison, WI, USA). Then pmirGLO-Wt/pmirGLO-Mt or miR-1299 mimics/miR-NCs were co-transfected into EC9706/KYSE30 cells. After 48 h of transfection, firefly luciferase activities of cells were measured via dual-luciferase reporter assays (Promega, Madison, WI, USA).

### RNA immunoprecipitation (RIP) assays

RIP assays were conducted to explore the relationship between lncRNA NEAT1 and miR-1299 via the Magna RIP RNA-Binding Protein Immunoprecipitation Kit (Millipore, MA, USA). Initially, digested EC9706 cells were harvested and lysed with complete RIP lysis buffer. Afterword, the supernatants of cell lysates were incubated with magnetic beads coated with anti-Ago 2 or control anti-IgG (Millipore, MA, USA) for 4 h at 4°C. Then, the immunoprecipitation complex was washed six times with cold RIP wash buffer. Finally, the RNA levels of extracted lncRNA NEAT1 and miR-1299 were detected by qRT-PCR.

### Statistical analysis

GraphPad Prism 5.0 (GraphPad Software, San Diego, USA) and Statistical SPSS 19.0 (IBM, NY, USA) software were used for statistical analysis. All assays were independently repeated three times under the same conditions. Moreover, all data were reported as the mean ± standard deviation (SD). Linear correlation analysis and independent sample t-tests were used for analysis. A *P*<0.05 was considered to indicate a statistically significant difference.

## Results

### lncRNA NEAT1 is upregulated in ESCC tissues and cell lines

We measured RNA levels of lncRNA NEAT1, miR-1299, and MMP2 in 32 ESCC tissues, as well as in multiple ESCC cell lines, via qRT-PCR. The results showed that lncRNA NEAT1 and MMP2 in ESCC tissues were significantly up-regulated compared with those in para-cancer tissues (*P<0*.*05*) (Figures 1A, 1C). Meanwhile, miR-1299 in ESCC tissues was significantly down-regulated (*P<0*.*05*) (Figure 1B). Analysis of clinicopathological characteristics indicated that miR-1299 levels in ESCC tissues were not significantly different as a function of gender, age, lymph-node metastasis status, or and TNM stage. In contrast, lncRNA NEAT1 and MMP2 in ESCC tissues with lymph-node metastasis were significantly higher than those without lymph-node metastasis (*P<0*.*05*); additionally, lncRNA NEAT1 and MMP2 in ESCC tissues at the TNM-III stage were significantly higher than those at stage I or stage II (*P<0*.*05*) (Table 2). Furthermore, lncRNA NEAT1 and MMP2 in three ESCC cell lines were significantly upregulated compared with those in normal human esophageal epithelial cells (Het-1A) (*P<0*.*05*) (Figures 1D, 1F). Conversely, miR-1299 levels in three ESCC cell lines were significantly downregulated compared with those in Het-1A cells (*P<0*.*05*) (Figure 1E). These results suggested that lncRNA NEAT1 and MMP2 were overexpressed in ESCC tissues and cells, whereas miR-1299 exhibited low expression. Collectively, these findings suggest that up-regulation of lncRNA NEAT1 was related to the progression of ESCC.

**Table 2.** Clinicopathologic characteristics and lncRNA NEAT1, MiR-1299, and MMP2 levels in 32 ESCC samples

| Characteristics | lncRNA NEAT1 levels | *P*-value | miR-1299 levels | *P*-value | MMP2 levels | *P*-value |
| --- | --- | --- | --- | --- | --- | --- |
| **Gender** |  |  |  |  |  |  |
| Female | 2.31±1.35 | 0.592 | 0.61±0.32 | 0.651 | 2.06±0.46 | 0.097 |
| Male | 2.56±1.26 |  | 0.66±0.37 |  | 2.33±0.43 |  |
| **Age** |  |  |  |  |  |  |
| ≤60 | 2.61±1.19 | 0.500 | 0.69±0.38 | 0.382 | 2.15±0.43 | 0.562 |
| >60 | 2.29±1.39 |  | 0.59±0.31 |  | 2.25±0.49 |  |
| **TNM Stage** |  |  |  |  |  |  |
| I + II | 1.97±1.18 | 0.016* | 0.70±0.35 | 0.233 | 2.02±0.38 | 0.008* |
| III | 3.05±1.20 |  | 0.55±0.32 |  | 2.44±0.45 |  |
| **Lymph-Node Metastasis** |  |  |  |  |  |  |
| Absence | 1.89±1.17 | 0.013* | 0.67±0.34 | 0.548 | 1.97±0.40 | 0.002* |
| Presence | 2.99±1.19 |  | 0.60±0.35 |  | 2.44±0.39 |  |

\*P<0.05

**Figure 1.**
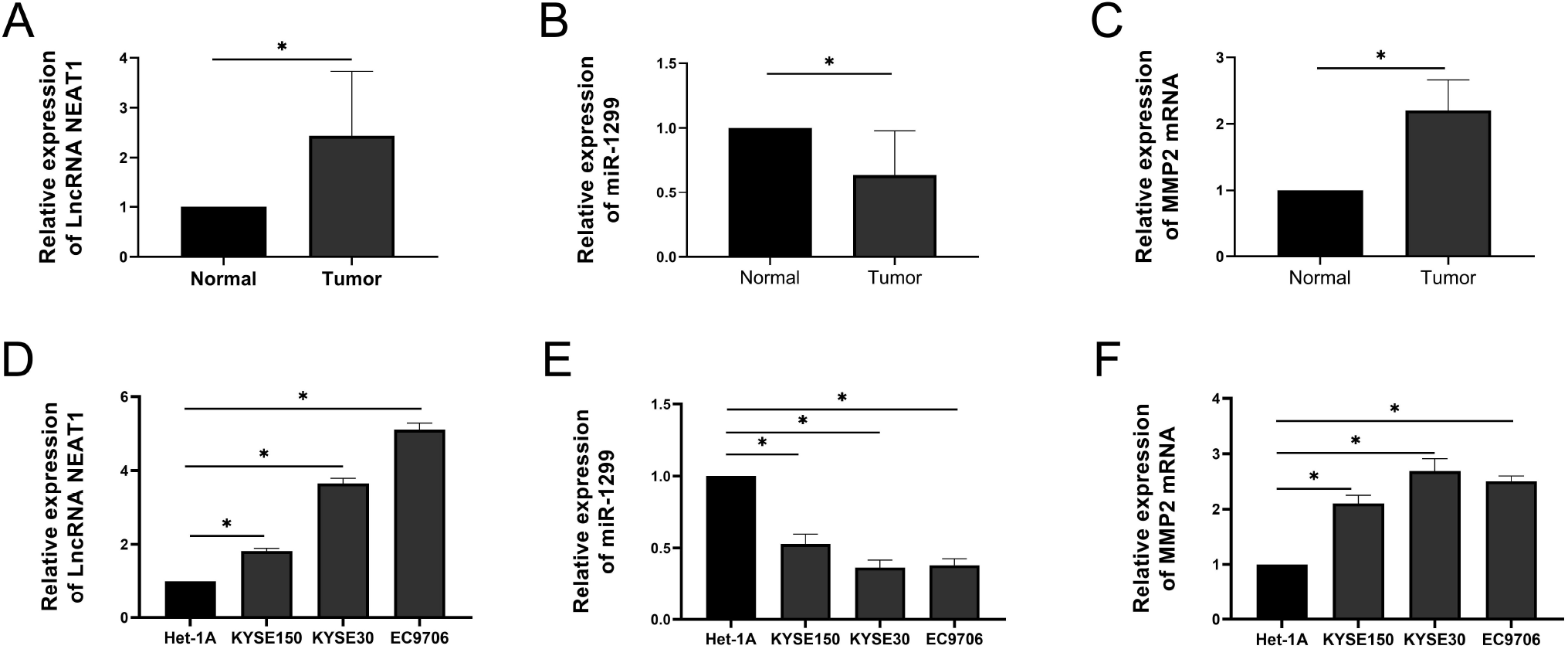
lncRNA NEAT1 is upregulated in ESCC tissues and cells. A) The expression levels of lncRNA NEAT1 in 32 ESCC tissues and their para-carcinoma tissues were analyzed by qRT-PCR. B) The expression levels of miR-1299 in 32 ESCC tissues and their para-carcinoma tissues were analyzed by qRT-PCR. C) The expression levels of MMP2 mRNA in 32 ESCC tissues and their para-carcinoma tissues were analyzed by qRT-PCR. D) Relative expression levels of lncRNA NEAT1 in Het-1A cells and ESCC cells were measured by qRT-PCR. E) Relative expression levels of miR-1299 in Het-1A cells and ESCC cells were measured by qRT-PCR. F) Relative expression levels of MMP2 mRNA in Het-1A cells and ESCC cells were measured by qRT-PCR. ^*^P<0.05

### lncRNA NEAT1 silencing impairs ESCC proliferation, invasion, and migration

KYSE30 and EC9706 cells were used to further investigate the function of lncRNA NEAT1. First, we verified the knockdown efficiency of lncRNA NEAT1 in KYSE30 and EC9706 cells (Figure 2A). Next, CCK8 assays were performed in KYSE30 and EC9706 cells. The results indicated that knockdown lncRNA NEAT1 significantly impaired cellular proliferation in KYSE30 and EC9706 cells (Figures 2B, 2C). Additionally, transwell assays demonstrated that lncRNA NEAT1 knockdown also attenuated invasion and migration of KYSE30 and EC9706 cells (Figures 2D, 2E). Therefore, these results validated that silencing of lncRNA NEAT1 inhibited proliferation, invasion, and migration of ESCC.

**Figure 2.**
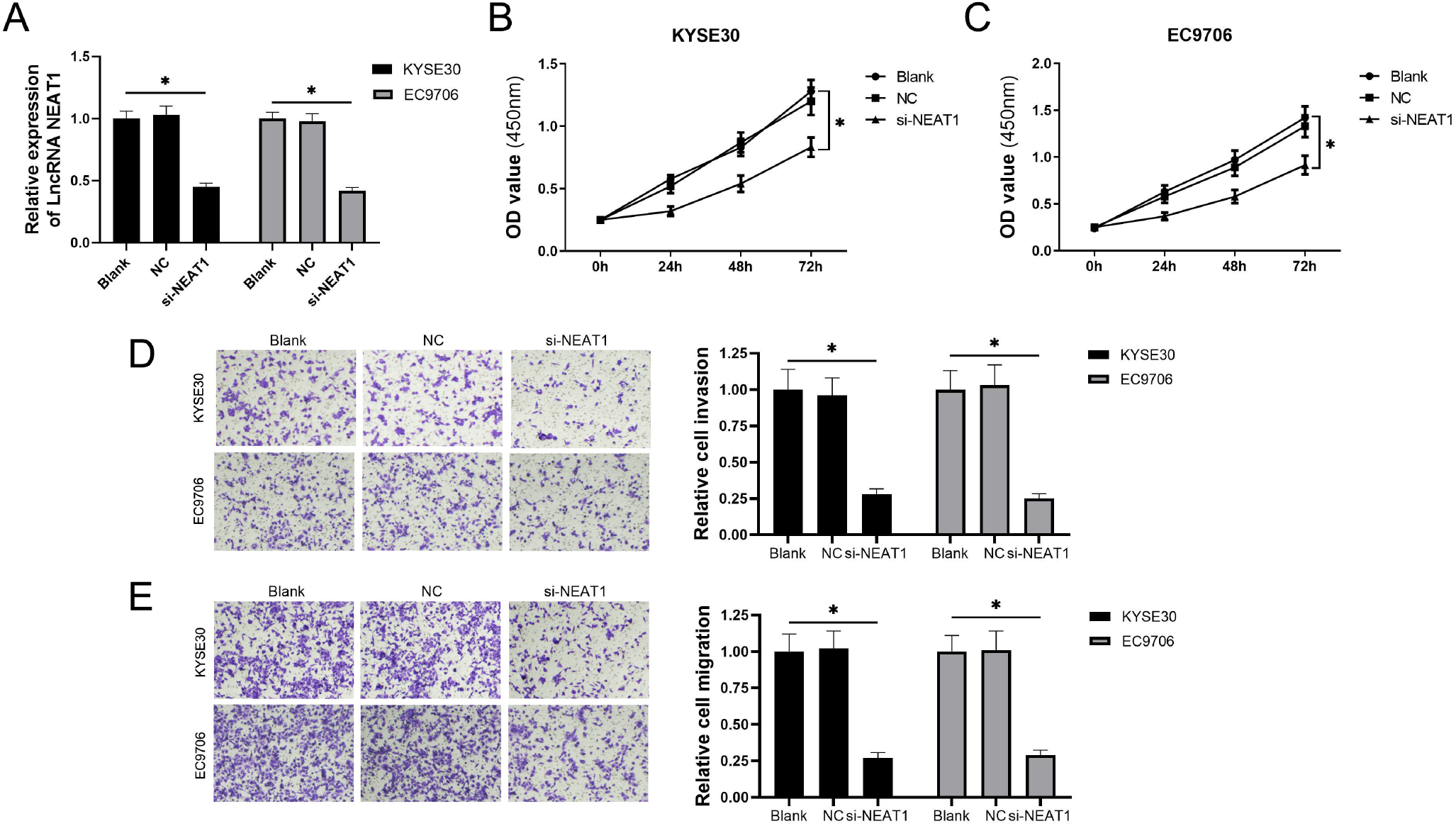
lncRNA NEAT1 silencing impairs ESCC proliferation, invasion, and migration. A) Relative expression levels of lncRNA NEAT1 in KYSE30 and EC9706 cells transfected with si-NC, or si-NEAT1. B) CCK8 assays revealed that si-NEAT1 impaired the proliferation in KYSE30 cells. C) CCK8 assays revealed that si-NEAT1 impaired the proliferation in EC9706 cells. D) Transwell assays of cell invasion revealed that lncRNA NEAT1 knockdown impaired invasion of KYSE30 and EC9706 cells. E) Transwell assays of cell migration revealed that lncRNA NEAT1 knockdown impaired migration of KYSE30 and EC9706 cells. ^*^P<0.05

### lncRNA NEAT1 sponges miR-1299 in ESCC cells

We next explored potential targets of lncRNA NEAT1 via bioinformatic analysis (LncBase v2 and miRDB), among which miR-1299 was found to have the highest target score (Figure 3A). Then, to verify the binding level of lncRNA NEAT1 with miR-1299, dual-luciferase reporter assays and RIP assays were executed with EC9706 cells. The data indicated that the activity of luciferase was inhibited by miR-1299 in pmirGLO-WT1,2,3 containing three binding sites (Figures 3B–3D). In addition, lncRNA NEAT1 and miR-1299 in anti-Ago2-immunoprecipitated complexes were significantly enriched compared with those in control anti-IgG-immunoprecipitated complexes (*P<0*.*05*) (Figure 3E), verifying that lncRNA NEAT1 and miR-1299 could simultaneously bind to anti-Ago2. The above data indicated that lncRNA NEAT1 interacted directly with miR-1299. Subsequently, to better elucidate the interaction of lncRNA NEAT1 and miR-1299 in ESCC, qRT-PCR assays were executed with KYSE30 and EC9706 cells. The data revealed that silencing of lncRNA NEAT1 induced up-regulation of miR-1299 in KYSE30 and EC9706 cells (Figure 3F). Additionally, we found that there was a significantly negative correlation between lncRNA NEAT1 and miR-1299 levels in ESCC tissues (Figure 3G). Taken together, these findings demonstrated that lncRNA NEAT1 sponges and negatively regulates miR-1299 as a ceRNA in ESCC cells.

**Figure 3.**
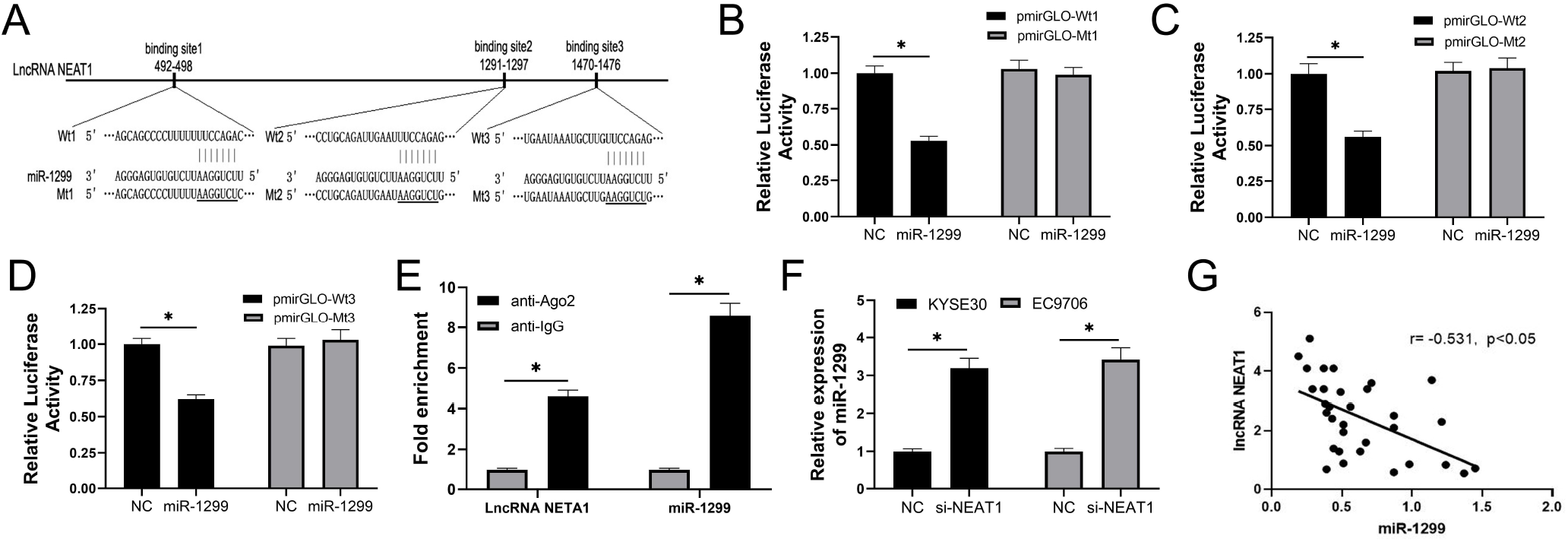
lncRNA NEAT1 sponges and negatively regulates miR-1299 in ESCC cells. A) Chart showing three inferred binding sites of lncRNA NEAT1 with miR-1299. The regions marked with an underscore were the mutated sites in lncRNA NEAT1. B) Dual-luciferase reporter assays were executed with EC9706 cells and showed that the activity of luciferase was inhibited by miR-1299 in pmirGLO-Wt1. C) Dual-luciferase reporter assays showed that the activity of luciferase was inhibited by miR-1299 in pmirGLO-Wt2. D) Dual-luciferase reporter assays showed that the activity of luciferase was inhibited by miR-1299 in pmirGLO-Wt3. E) RIP assays were executed with EC9706 cells and showed that lncRNA NEAT1 and miR-1299 in anti-Ago2-immunoprecipitations were enhanced compared with those in control anti-IgG-immunoprecipitations. F) Silencing of lncRNA NEAT1 promoted miR-1299 expression in KYSE30 and EC9706 cells. G) A significant negative linear correlation was found between lncRNA NEAT1 levels and miR-1299 levels in ESCC tissues. ^*^P<0.05

### MiR-1299 suppresses invasion and migration by targeting MMP2 in ESCC

First, we confirmed that increased miR-1299 negatively regulated the expression of MMP2 at the mRNA level (Figure 4A). Second, we substantiated the suppressive effect of miR-1299 acting on invasion and migration in KYSE30 and EC9706 cells. Moreover, we discovered that the suppressive effect of miR-1299 could be reversed by pcDNA-MMP2 (Figures 4B, 4C). Finally, by performing dual-luciferase reporter assays, we validated that miR-1299 targeted MMP2 with the predicted binding sites (Figures 4D–4F). In summary, these results demonstrated that miR-1299 directly targeted MMP2 and weakened its mediated invasion and migration in KYSE30 and EC9706 cells.

**Figure 4.**
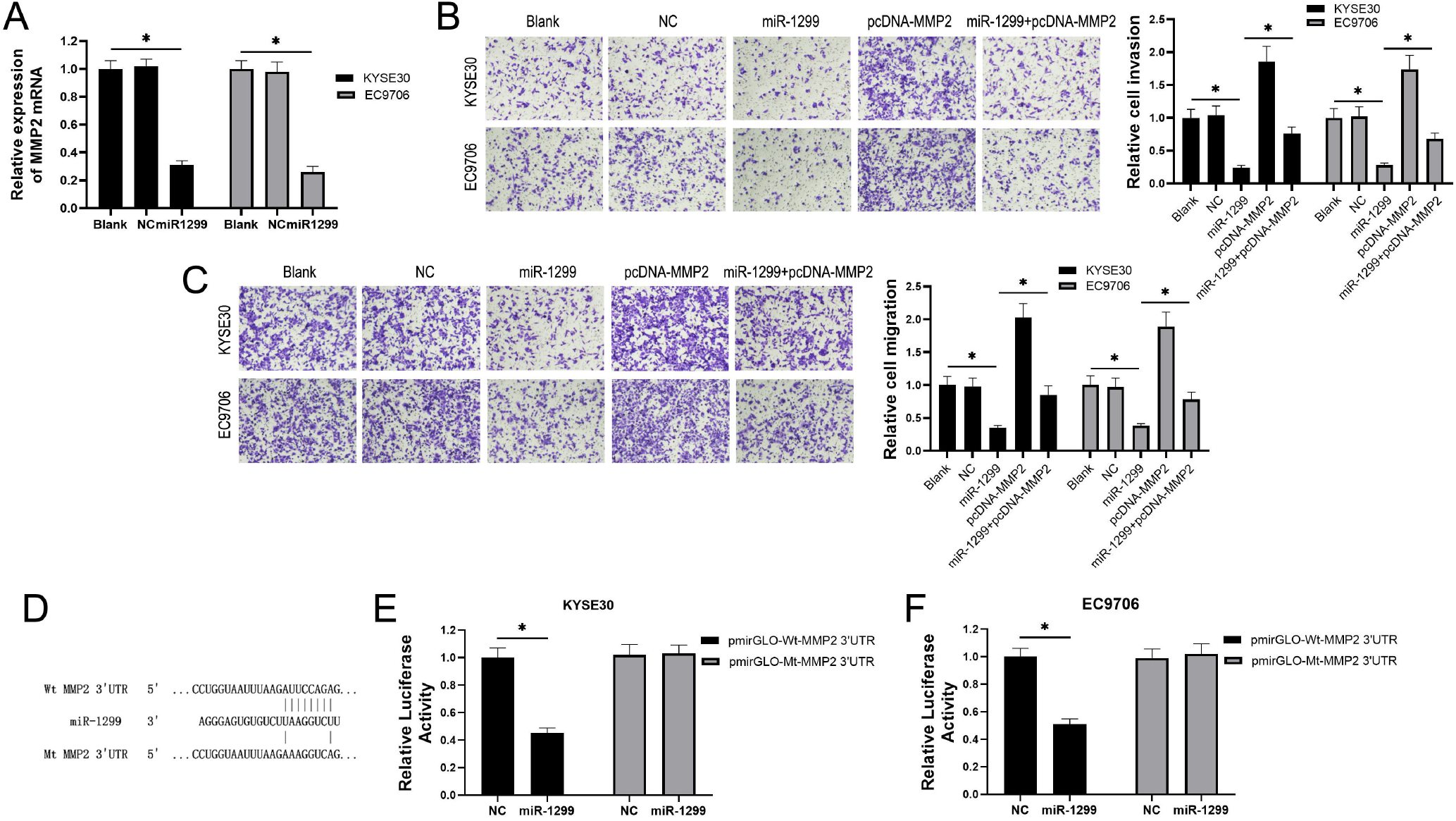
MiR-1299 weakens invasion and migration through targeting MMP2 in ESCC. A) The expression levels of MMP2 mRNA in KYSE30 and EC9706 cells transfected with miR-NC and miR-1299 mimics were analyzed by qRT-PCR. B) Transwell assays of cell invasion in KYSE30 and EC9706 cells transfected with miR-NC, miR-1299 mimics, pcDNA-MMP2, and miR-1299 mimics+pcDNA-MMP2. C) Transwell assays of cell migration in KYSE30 and EC9706 cells transfected with miR-NC, miR-1299 mimics, pcDNA-MMP2, and miR-1299 mimics+pcDNA-MMP2. D) Chart of predicted miR-1299 binding sites of MMP2 3’-UTR. E) MiR-1299 inhibited the activity of luciferase in KYSE30 cells co-transfected with pmirGLO-Wt-MMP2 3’-UTR. F) MiR-1299 inhibited the activity of luciferase in EC9706 cells co-transfected with pmirGLO-Wt-MMP2 3’-UTR. ^*^P<0.05

### lncRNA NEAT1 silencing suppresses MMP2-mediated invasion by reducing the competitively-binding miR-1299 in ESCC cells

Next, we investigated whether lncRNA NEAT1 regulated ESCC invasion by miR-1299/MMP2. Transwell rescue assays were performed with KYSE30 and EC9706 cells. As shown in Figures 5A and 5B, silencing of lncRNA NEAT1 suppressed invasion in KYSE30 and EC9706 cells. Meanwhile, miR-1299 inhibitors and pcDNA-MMP2 promoted invasion. The data validated that miR-1299 inhibitors and pcDNA-MMP2 significantly rescued the invasion of KYSE30 and EC9706 cells following silencing of lncRNA NEAT1 (*P*<0.05). Thus, our findings elucidated that silencing of lncRNA NEAT1 reduced the competitively-binding miR-1299 to inhibit MMP2-mediated invasion in ESCC cells.

**Figure 5.**
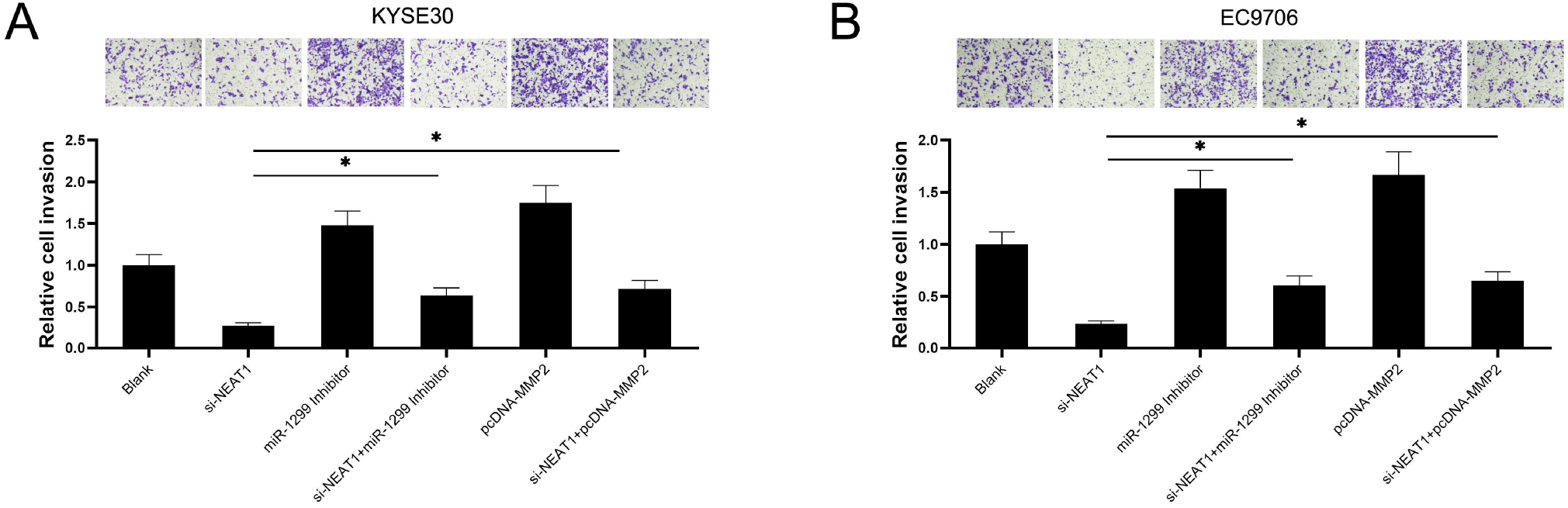
lncRNA NEAT1 silencing suppresses MMP2-mediated invasion by reducing the competitively-binding miR-1299 in ESCC cells. A) Transwell rescue assays of cell invasion in KYSE30 cells transfected with si-NEAT1, miR-1299 inhibitor, si-NEAT1+miR-1299 inhibitor, pcDNA-MMP2, and si-NEAT1+pcDNA-MMP2. B) Transwell rescue assays of cell invasion in EC9706 cells transfected with si-NEAT1, miR-1299 inhibitor, si-NEAT1+miR-1299 inhibitor, pcDNA-MMP2, and si-NEAT1+pcDNA-MMP2. ^*^P<0.05

## Discussion

Many studies have demonstrated that lncRNAs perform crucial roles in the progression of malignant tumors and may also represent novel targets for oncotherapy[27]. Therefore, the current study was performed to investigate the function of lncRNA NEAT1 in ESCC. We found that lncRNA NEAT1 was highly expressed in ESCC tissue samples and served as a ceRNA via sponging miR-1299 to upregulate MMP2, which ultimately promoted the progression of ESCC.

Initially, in our present study, we discovered the lncRNA NEAT1 was upregulated in 32 ESCC tissue samples and in three different cell lines. Furthermore, the expression level of lncRNA NEAT1 was closely associated with lymph-node metastasis and clinical stage. Additionally, silencing of lncRNA NEAT1 suppressed cell proliferation, invasion, and migration, consistent with previous research.[28] Moreover, we found that lncRNA NEAT1 sponged and inhibited miR-1299. Recently, lncRNAs have been confirmed to serve as ceRNAs via interacting with miRNA seed sequences[9, 10, 12]. For instance, lncRNA MEG3 participates in the inhibition of cell proliferation and invasion by serving as a ceRNA to regulate the expression of FOXO1 and E-cadherin via competitively binding miR-9 in ESCC[29]. In addition, lncRNA HOTAIR promotes the progression of EMT by acting as a ceRNA to regulate ZEB1 expression through the sponging of miR-130a-5p in ESCC[30]. Therefore, we hypothesized that lncRNA NEAT1 might competitively bind specific miRNAs to exert the inhibitory effects in ESCC. To test this hypothesis, we searched for potential miRNAs interacting with lncRNA NEAT1 through miRDB and LncBase v2. We ultimately identified that miR-1299 had the highest target score with lncRNA NEAT1 and exhibited binding to three sites simultaneously. Moreover, qRT-PCR assays suggested that the expression of miR-1299 was negatively correlated with that of lncRNA NEAT1 in ESCC cells and tissue samples. Furthermore, based on dual-luciferase reporter assays and RIP assays, we verified that miR-1299 was a direct target of lncRNA NEAT1 and possessed three binding regions. Interestingly, miR-1299 has been confirmed to be a tumor-inhibiting factor[21-25].Collectively, these results demonstrate that lncRNA NEAT1 acted as a ceRNA to enhance cell proliferation, invasion, and migration by sponging miR-1299 in ESCC.

Subsequently, we found that miR-1299 overexpression reduced the expression of MMP2 mRNA. Previous studies have shown that miR-1299 inhibits the progression of diverse human tumors by targeting mRNAs for molecular proteins[25, 31]. For example, miR-1299 inhibits the growth, invasion, and metastasis of prostate cancer via regulating NEK2[22]. Inhibition of miR-1299 expression increases the proliferation, cell-cycle progression, and invasion of HCC cells by promoting CTNND1 expression[32]. In the current study, we found that miR-1299 negatively regulated MMP2 expression and the pcDNA-MMP2 weakened the ability of miR-1299 to inhibit invasion and migration in KYSE30 and EC9706 cells, suggesting that miR-1299 was involved in MMP2-mediated invasion and migration. Furthermore, bioinformatics and dual-luciferase assays confirmed that MMP2 was a direct target of miR-1299 in ESCC cells. Finally, we found that silencing of lncRNA NEAT1 also inhibited MMP2-mediated invasion by reducing the competitively binding miR-1299 in KYSE30 and EC9706 cells. By matching the seed sequences of lncRNA NEAT1 and MMP2 binding with miR-1299, we discovered that lncRNA NEAT1 and MMP2 shared the same base sequences for miR-1299. Moreover, based on transwell rescue assays, si-NEAT1 suppressed invasion, whereas miR-1299 inhibitors and pcDNA-MMP2 markedly reversed the suppressive effect of silencing lncRNA NEAT1 in KYSE30 and EC9706 cells. These results indicate that there is an ESCC-specific ceRNA network based on expression profiles of lncRNAs, miRNAs, and mRNAs[33]. Furthermore, similar ceRNA networks have been elucidated in numerous previous studies[10,12,14,16]. Collectively, these results strongly suggest that the supressed effect of lncRNA NEAT1 silencing on MMP2 in our present study was accomplished via the miR-1299 pathway.

In conclusion, we found that lncRNA NEAT1 was overexpressed in ESCC tissues and cells, and was associated with lymph-node metastasis and TNM stage.Silencing of lncRNA NEAT1 inhibited ESCC proliferation, invasion, and migration via reducing the competitively-binding miR-1299 and enhancing miR-1299-targeted suppression of MMP2. Our present findings provide novel evidence that lncRNAs function as ceRNAs of miRNAs, and that lncRNA NEAT1 may represent a promising target for ESCC therapy and rehabilitation.

## Acknowledgments

Not applicable.

